# Murine monoclonal antibodies against RBD of SARS-CoV-2 neutralize authentic wild type SARS-CoV-2 as well as B.1.1.7 and B.1.351 viruses and protect *in vivo* in a mouse model in a neutralization dependent manner

**DOI:** 10.1101/2021.04.05.438547

**Authors:** Fatima Amanat, Shirin Strohmeier, Wen-Hsin Lee, Sandhya Bangaru, Andrew B. Ward, Lynda Coughlan, Florian Krammer

## Abstract

After first emerging in December 2019 in China, severe acute respiratory syndrome 2 (SARS-CoV-2) has since caused a pandemic leading to millions of infections and deaths worldwide. Vaccines have been developed and authorized but supply of these vaccines is currently limited. With new variants of the virus now emerging and spreading globally, it is essential to develop therapeutics that are broadly protective and bind conserved epitopes in the receptor binding domain (RBD) or the whole spike of SARS-CoV-2. In this study, we have generated mouse monoclonal antibodies (mAbs) against different epitopes on the RBD and assessed binding and neutralization against authentic SARS-CoV-2. We have demonstrated that antibodies with neutralizing activity, but not non-neutralizing antibodies, lower viral titers in the lungs when administered in a prophylactic setting *in vivo* in a mouse challenge model. In addition, most of the mAbs cross-neutralize the B.1.351 as well as the B.1.1.7 variants *in vitro*.

**Importance:** Crossneutralization of SARS-CoV-2 variants by RBD-targeting antibodies is still not well understood and very little is known about the potential protective effect of non-neutralizing antibodies *in vivo*. Using a panel of mouse monoclonal antibodies, we investigate both of these aspects.

## Introduction

SARS-CoV-2 first emerged in late 2019 in the province of Hubei in China, spread rapidly throughout the globe, and has since caused the ongoing coronavirus disease 2019 (COVID-19) pandemic (1, 2). Millions of infections have occurred globally and over two million deaths have been caused by this novel coronavirus. Over a hundred vaccines are currently in clinical development with three vaccines authorized for use in humans under the emergency use authorization (EUA) in the United States by the Food and Drug Administration (FDA) and several additional ones approved in Europe, Latin America and Asia. Furthermore, several monoclonal antibodies (mAbs) as cocktails and an antiviral, remdesivir, have been authorized for use in humans as therapeutics and numerous others including antivirals are in development (3–6).

SARS-CoV-2, a positive-sense single-stranded RNA virus from the *Coronaviridae* family, is closely related to SARS-CoV-1 which caused a major outbreak in 2002-2004. Both viruses use the same receptor for entry into host cells, human angiotensin converting enzyme (hACE2) (7, 8). The receptor binding domain (RBD) which is part of the spike protein of the virus can bind to hACE2 and mediate entry and thus, the spike protein makes for an excellent target for vaccines and therapeutics (9). It has been observed that serum from infected individuals as well as from vaccinated individuals contains a high level of antibodies against the spike protein and this serum shows high neutralizing activity (10–12). Antibodies induced by natural infection with SARS-CoV-2 correlate with protection and vaccination has been shown to be highly efficacious and effective as well. However, it is still crucial to develop therapeutics that can be used to treat individuals that are infected with SARS-CoV-2, particularly those at high risk for severe disease. While mAbs have been developed and approved for use, there remains a significant concern from the virus acquiring mutations that would lead to escape rendering the mAbs and vaccines inefficient in blocking virus and stopping replication of the virus in the body. Several lineages of SARS-CoV-2 with distinct and sometimes additional mutations in the spike protein have emerged over the last year)(13). Mutations in the RBD region of the spike protein are a serious concern as most neutralizing antibodies target the RBD and block entry. Another region heavily mutated in the new circulating variant viruses is the N-terminal domain (NTD) which is also a target of neutralizing antibodies(14). Hence, the efficacy of vaccines and therapeutics could be compromised as more and more mutations in the NTD and RBD occur and persist in nature (15, 16).

In this study, we isolated and characterized fourteen mouse mAbs against the RBD of SARS-CoV-2 and assessed their binding to recombinant RBD and spike protein as well as tested their ability to neutralize live virus. In addition, we tested if non-neutralizing mAbs can lower viral loads in a mouse challenge model. Due to the new variants of concern which have been detected, we also tested if mAbs can bind mutant RBDs that contain single amino acid changes as well as multiple mutations found in B.1.1.7, B.1.351 and P.1 RBDs. Lastly, we tested our panel of neutralizing mAbs against the B.1.1.7 virus isolate as well as B.1.351 virus isolate.

## Results

### Generation of monoclonal antibodies

After two vaccinations of BALB/c mice with recombinant RBD protein supplemented with poly I:C, murine hybridoma technology was used to generate hybridoma cell lines that secreted RBD-specific monoclonal antibodies (17–19). Fourteen unique hybridoma lines were isolated and picked that produced IgGs (**Table 1**). Twelve monoclonal antibodies belonged to the IgG1 isotype while two monoclonal antibodies were from the IgG2a subclass.

**Table 1.**

| mAb | Isotype |
| --- | --- |
| KL-S-1B5 | IgG1 |
| KL-S-1D2 | IgG2a |
| KL-S-1D11 | IgG1 |
| KL-S-1E10 | IgG1 |
| KL-S-1F7 | IgG1 |
| KL-S-1H12 | IgG1 |
| KL-S-2A1 | IgG1 |
| KL-S-2A5 | IgG1 |
| KL-S-2C3 | IgG2a |
| KL-S-2F1 | IgG1 |
| KL-S-2F9 | IgG1 |
| KL-S-2G9 | IgG1 |
| KL-S-3A5 | IgG1 |
| KL-S-3A7 | IgG1 |

### All antibodies bind the RBD of SARS-CoV-2 and six mAbs can neutralize live virus

Once all antibodies were purified from hybridoma supernatant, a standard enzyme-linked immunosorbent assay (ELISA) was performed to assess binding of the monoclonal antibodies to the RBD of SARS-CoV-2 (**Figure 1A**), full spike of SARS-CoV-2 (**Figure 1B**), and RBD of SARS-CoV-1 (**Figure 1C**). All antibodies bound well to SARS-CoV-2 RBD and most had very low minimal binding concentrations (MBC). Of note, the MBC values for KL-S-1B5 and KL-S-2A1 (0.1 ug/ml) against SARS-CoV-2 RBD were higher than the rest of the antibodies, indicating lower affinity. Next, antibodies were tested in an ELISA against the full spike protein of SARS-CoV-2 (**Figure 1B**). It is interesting to note that while most mAbs bound well with low MBC values, KL-S-1B5, KL-S-1D11 and KL-S-2A1 had higher MBC values against spike compared to the RBD of SARS-CoV-2. It is possible that the epitope of these antibodies is partially occluded on the full spike compared to the RBD protein when expressed alone. To determine if antibodies were cross-reactive to the RBD of SARS-CoV-1, an ELISA was performed (**Figure 1C**). Most mAbs could not bind the RBD of SARS-CoV-1 except KL-S-1E10, KL-S-2A5, KL-S-3G9, and KL-S-3A5. To assess the functionality of the mAbs, all fourteen mAbs were tested in a microneutralization assay with authentic SARS-CoV-2 for their ability to neutralize live virus (**Figure 1D**). Six mAbs (43%) neutralized live virus well with low IC_50_ values (0.1-1 ug/ml) indicating that low concentrations are capable of blocking virus entry and/or replication. Notably, KL-S-1D2 and KL-S-3A7 are extraordinarily low IC_50_ values lower than 1 ug/ml.

**Figure 1.**
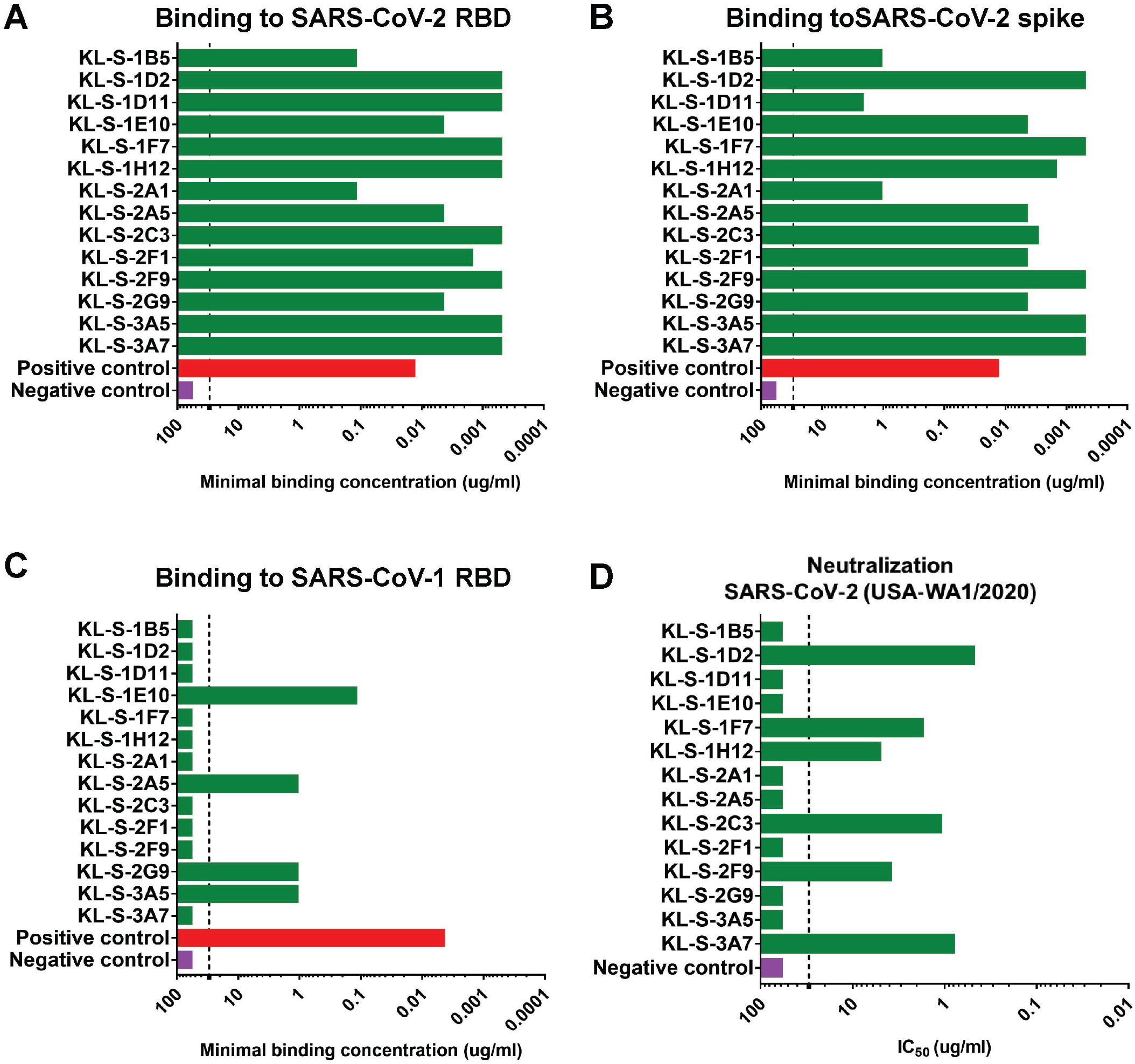
All mAbs bind to recombinant RBD and six mAbs neutralize SARS-COV-2. **(A)** Binding of all isolated mAbs (n=14) via an ELISA assay was assessed against recombinant RBD protein and the MBC values are shown. The positive control used was a human antibody against SARS-CoV-1 RBD, CR3022 while the negative control used was a mouse anti-influenza H10 antibody. Binding of all isolated mAbs was also tested on ELISA against the spike protein of SARS-CoV-2 **(B)** as well as SARS-CoV-1 RBD **(C)**. **(D)** Neutralization activity of all mAbs was tested in a microneutralization assay with authentic SARS-CoV-2 (USA-WA1/2020) starting at 30 ug/ml and testing subsequent 3-fold dilutions. The cells were stained for nucleoprotein of SARS-CoV-2 and the IC_50_ values were calculated via non-linear regression fit. All experiments were performed with duplicates.

### Antibodies can lower viral titers *in vivo* in a mouse challenge model

To further study the biological functionality of these mAbs, all mAbs were tested *in vivo*. Hence, an animal model was utilized to test if antibodies are able to block viral entry and thus lower titers in the lung. Since mouse ACE2 does not facilitate entry of SARS-CoV-2, an adenovirus that expresses the human ACE2 gene was used to first transduce mice (20, 21). Five days later, monoclonal antibodies were administered at 10 mg/kg two hours prior to infection with SARS-CoV-2 and lungs were collected on day 3 and day 5 post infection to assess viral titers via a plaque assay. Only neutralizing mAbs were able to confer a protective benefit and lowered viral titers in the lungs (**Figure 2A-B**). On day 3, the group that received KL-S-1D2 and KL-S-2C3 had significantly less virus in the lungs compared to other groups highlighting the antibodies’ ability to protect *in vivo*. KL-S-1F7, KL-S-1H12 and KL-S-2F9 treated animals had approximately two logs less virus in their lung compared to the negative control on day 3 (**Figure 2A**). The negative control used here was an irrelevant purified antibody, binding to influenza virus H10 hemagglutinin (18). On day 5, groups that received the six neutralizing mAbs had little to no virus in their lungs (**Figure 2B**). One mouse in the KL-S-2C3 group and one mouse in the KL-S-2F9 showed residual virus in the lungs which could be a caveat of animal model used. None of the non-neutralizing antibodies conferred any protective benefit and all of these mAbs belonged to the IgG1 isotype.

**Figure 2.**
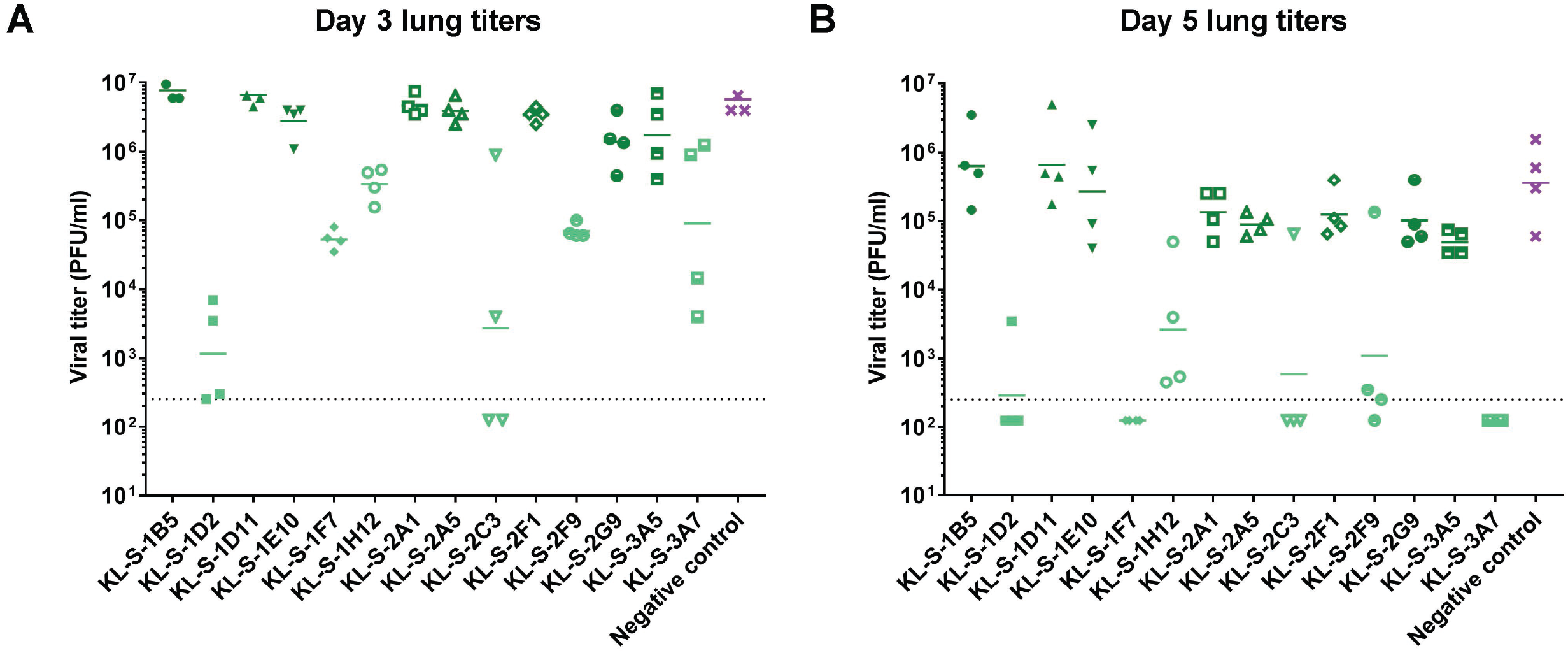
Only neutralizing mAbs lower viral loads *in vivo* in a AdV-hACE2 mouse challenge model. The protective efficacy of the mAbs was assessed *in vivo* in a prophylactic setting. Mice were administered 2.5* 10^8^ PFUs per mouse of AdV-hACE2 and five days later, mice were administered each mAb (n=4) at 10 mg/kg and challenged with 10^5^ PFUs of SARS-CoV-2. Viral titers in the lungs were assessed at day 3 **(A)** and day 5 **(B)** via a plaque assay. Mice in the negative control group received a mouse anti-influenza H10 antibody.

### Neutralizing antibodies eliminate viral presence in the lungs and little differences were found between the groups in terms of lung pathology

In addition to assessment of viral titers in the lungs in a prophylactic setting, we also wanted to test if mAbs can protect from inflammation and/or tissue damage in the lungs or lead to enhanced disease which has been noted for SARS-CoV-1 (22). Lungs were harvested on day 4 post vaccination from all the antibody groups (n=2) and subjected to pathological analysis (Histowiz) such as hematoxylin and eosin (H&E) staining as well as immunohistochemistry (IHC) using an antibody specific for the nucleoprotein of SARS-CoV-2. A 5-point grading scheme that took into account six different parameters (“perivascular inflammation”, “bronchial/bronchiolar epithelial degeneration/necrosis”, “bronchial/bronchiolar inflammation”, “intraluminal debris”, “alveolar inflammation” and “congestion/edema”) was utilized to score lung sections. Interestingly, mice from all groups treated with antibodies displayed some pulmonary histopathological lesions of interstitial pneumonia (**Figure 3**). This could be a result of the high dose (10^5^ PFU per mouse) of SARS-CoV-2 used. The group that received only the AdV-hACE2 exhibited some microscopic lesions of perivascular, peri-bronchiolar and alveolar inflammation and had much lower scores compared to the antibody groups that received AdV-hACE2 plus SARS-CoV-2 (**Supplementary figure 1**). This demonstrates that there is some mild inflammation associated with intranasal administration of AdV-hACE2 alone and this has been observed in earlier studies (20). Histopathological lesions were uniformly absent in the mock group that received no treatment and were basically just naïve mice. Clinical scores were slightly higher for groups KL-S-1E10, KL-S-2A5 and KL-S-3A5. Both of these antibodies are non-neutralizing, but this could be a result of external variables in the experimental setup.

**Figure 3.**
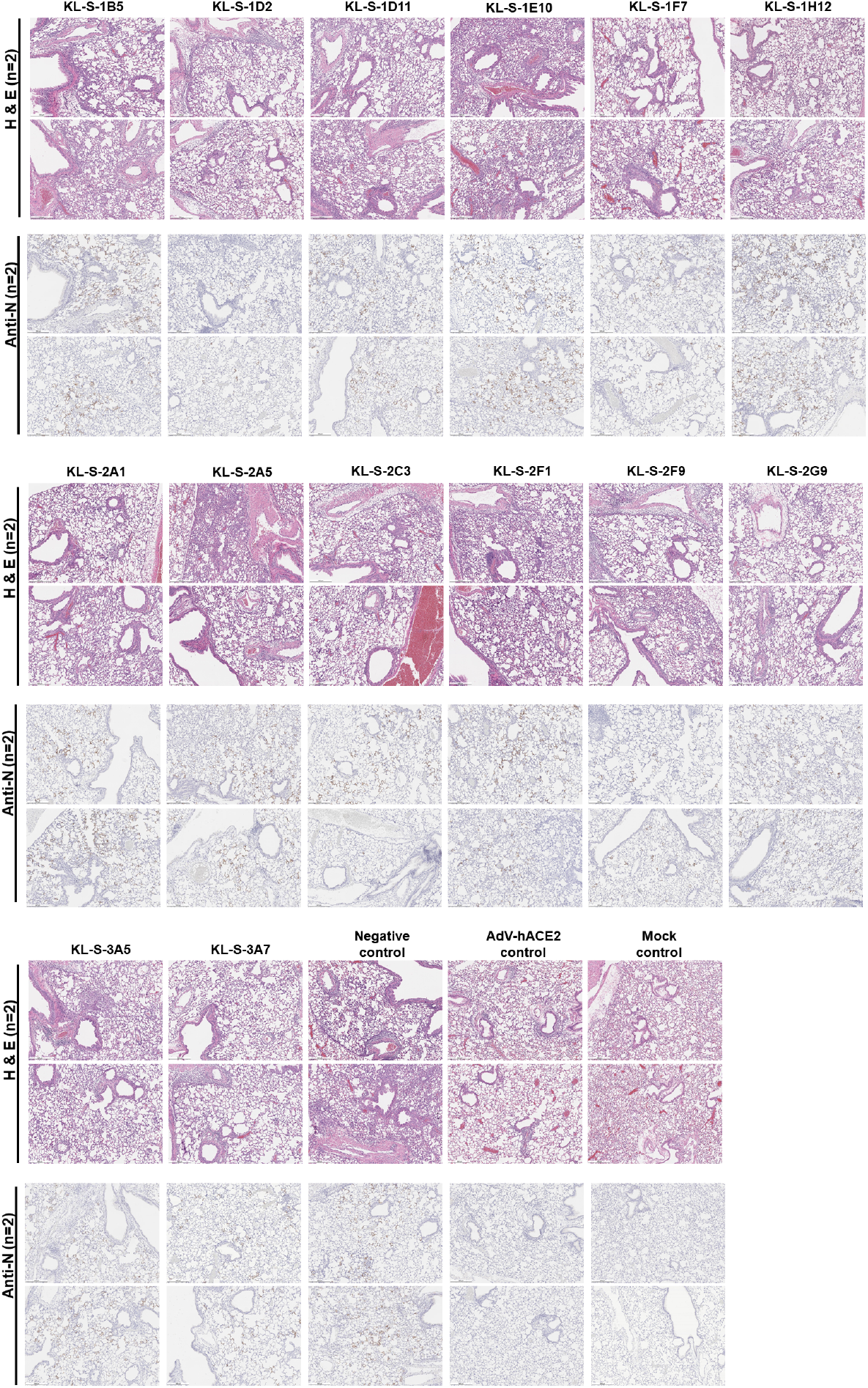
Immunopathological effects post mAb administration in the lungs. **(A)** In order to assess if antibodies can have any negative immunopathological effects, lungs were harvested from each antibody group (n=2) as shown. Two mice received only the AdV-hACE2 while two mice were naïve. An anti-nucleoprotein antibody was used to check for presence of virus in the lungs. **(B)**

In terms of the nucleoprotein staining via IHC, it is clear that all neutralizing mAbs except KL-S-1H12 blocked entry and thus, very little staining for nucleoprotein was observed on day 4. This correlates well with the lung titers found in figure 2 as antibodies blocked viral entry/replication and lowered viral load in the lungs.

### Antibodies maintain binding to most variant RBDs

Several variants of concern (VOC) with mutations in their RBD are circulating. In addition, studies on mAbs escape, in vitro evolution of the spike and clinical isolates from immunosuppressed patients have reported a variety of single mutations that may influence antibody binding to RBD. We expressed a number of these RBDs including N439K, Y453F, E484K, N501Y (B.1.1.7) and the RBDs of B.1.351 and P.1 and performed ELISAs on them using our mAbs. Such analysis can demonstrate the epitope of the antibody or the single amino acid that is crucial in its native state for binding. The neutralizing mAbs and non-neutralizing mAbs are shown separately (**Figure 4A and Figure 4B**) KL-S-1D2 maintained binding to all RBDs but lost complete binding to K487R RBD (**Figure 4A**). This could be a very crucial amino acid for the antibody to maintain its footprint on the RBD. KL-S-2C3 bound at only 30% to the N487R RBD compared to wild type RBD. KL-S-1H12 lost a lot of binding to E484K, F486A, F490K and the B.1.351 RBD (**Figure 4A**). KL-S-1D11, KL-S-1F7, KL-S-2F9, KL-S-2G9 and KL-S-3A7 were able to bind all mutant RBDs at a level of 50% or even more compared to wild type RBD (**Figure 4A**). KL-S-1E10 and KL-S-2A1 lost binding to a large number of mutant RBDs (**Figure 4B**). The ability to bind all RBDs could be a function of antibody affinity which, when high, can allow the antibody to maintain its footprint. In general, neutralizing mAbs had comparable binding to both wild type and most mutant RBDs. To ensure that the ELISA setup was comparable, an anti-histidine antibody was used as a positive control.

**Figure 4.**
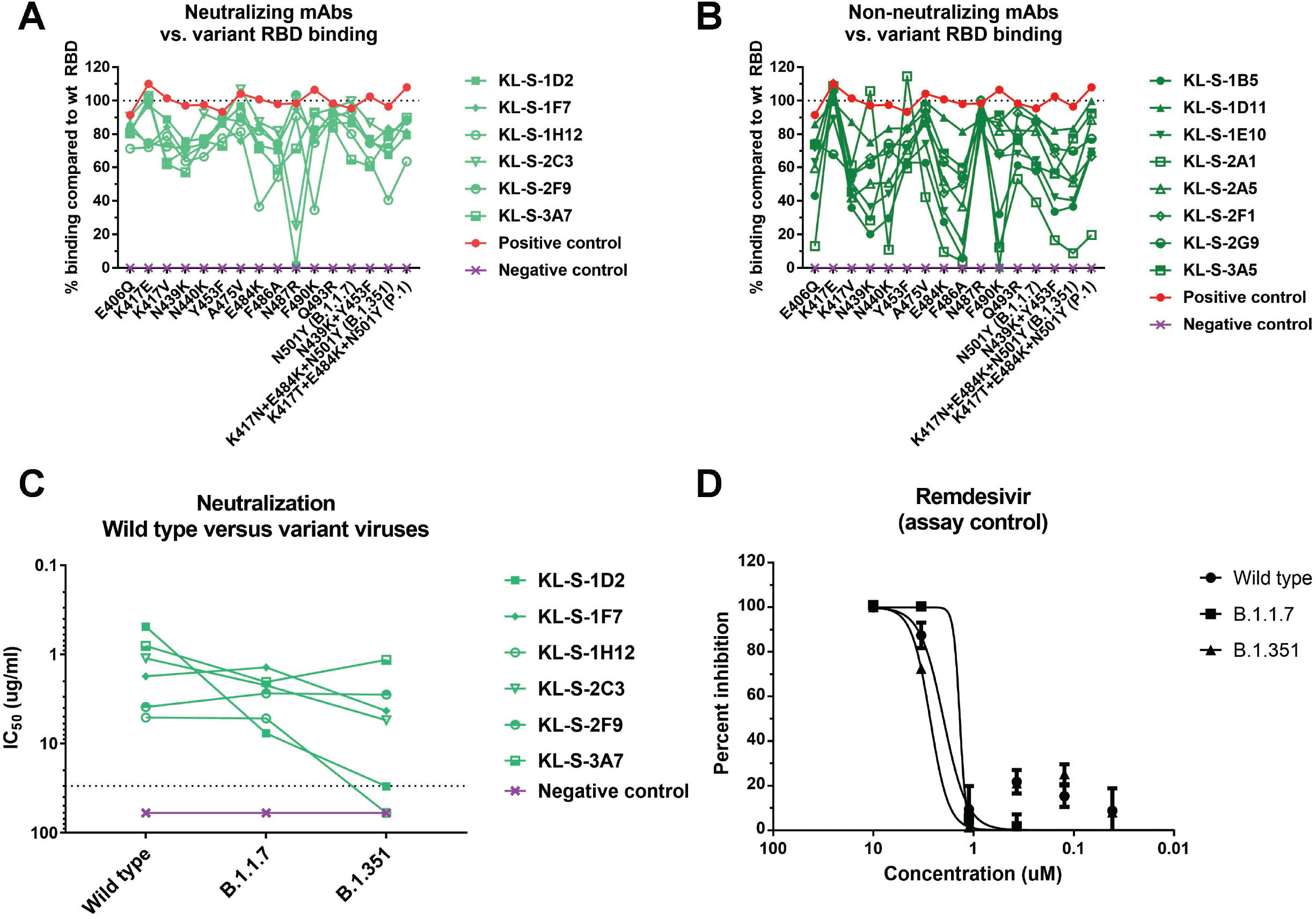
Binding of mAbs to variant RBDs as well cross-neutralization of B.1.1.7 and B.1.351 variant viruses. All fourteen antibodies (**A-B**) were tested in ELISA assays for binding to RBDs that contain single or multiple mutations found in new variants. The line at 100% indicates binding to wild type and binding to each mutant RBD is graphed as percent binding compared to wild type. A negative control mAb, anti-influenza H10, was run against all the RBDs to ensure that there is no unspecific binding. A positive control, anti-histidine antibody, was used to ensure that the RBD proteins that have a hexa-histidine tag are coated properly. Neutralizing mAbs **(A)** and non-neutralizing mAbs **(B)** are shown separately. **(C)** A microneutralization assay was performed to test whether the neutralizing mAbs can also neutralize new variant viruses, B.1.1.7 and B.1.351. IC_50_ values of the six neutralizing mAbs for each virus are shown. **(D)** Remdesivir was also run on a neutralization assay against the wild type virus, B.1.1.7 isolate as well as B.1.351 isolate.

### Four mAbs maintain neutralizing activity to B.1.351 virus while all six mAbs neutralize B.1.1.7 virus

Since binding may not be directly related to neutralization, we wanted to assess if antibodies that were generated by vaccination of mice with wild type RBD can neutralize new variant viruses. These variant viruses carry mutations in the RBD and can escape neutralization by monoclonal antibodies easily if their native epitope has been disrupted (6, 16). Notably, all antibodies that neutralized wild type SARS-CoV-2 were also able to neutralize the B.1.1.7 virus and this is not surprising as the only mutation present in the RBD of this virus is N501Y (**Figure 4C**). However, KL-S-1D2 and KL-S-1H12 completely lost neutralizing activity towards B.1.351 virus while the remaining four neutralizing mAbs maintained activity to this virus, although to various degrees (**Figure 4C**). KL-S-1D2 bound the RBD of B.1.351 at around 70% compared to wild type but loss of neutralization could be due to the epitope being presented differently on the full spike compared to RBD alone, leading to a loss of affinity. KL-S-1H12 showed much lower binding to E484K RBD as well as the B.1.351 RBD and this lower binding capability might be the reason for the loss of neutralization. The IC50 value for KL-S-1F7 was 1.7 ug/ml for the wild type virus, 1.4 ug/ml for the B.1.1.7 virus and 4.3 ug/ml for the B.1.351 virus. The IC50 value for KL-S-2C3 was 1.1 ug/ml for the wild type virus, 2.2 for the B.1.1.7 virus, and 5.5 ug/ml for the B.1.351 virus (**Figure 4C**).

### Three antibodies block ACE2 from binding the RBD

To study where the neutralizing antibodies bind on the RBD, structural analysis was performed, and negative stain three dimensional reconstructions were obtained for four of the neutralizing antibodies (**Figure 5**). KL-S-1H12 and KL-S-2F9 did not form stable complexes and were therefore not amenable to image analysis, which could be a result of pH changes or low affinity/avidity of the Fab. KL-S-2C3 (**Figure 5A**), KL-S-1D2 (**Figure 5B**), and KL-S-3A7 (**Figure 5C**) overlap with the ACE2 binding site, consistent with blockade of ACE2 binding to the RBD. The antibodies approach at different angles and appear to belong to Class 1 (KL-S-3A7 and KL-S-1D2) and Class 2 (KL-S-2C3) RBD epitopes (23). KLS-1F7 binds lower on the RBD to a similar epitope as S309 (**Figure 5D**) (24).

**Figure 5.**
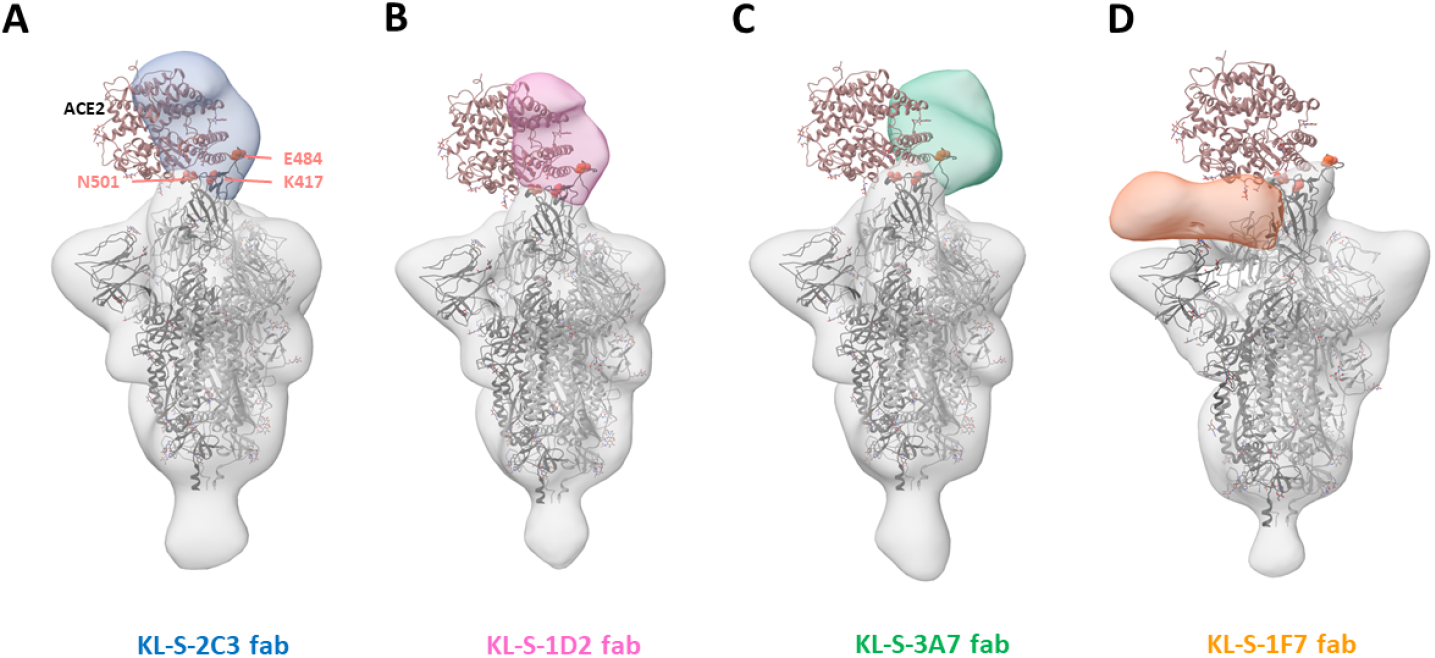
Negative stain EM analysis of Fabs bound to SARS-CoV-2 S trimer. 3D reconstructions of Fabs (**A**) KL-S-2C3 (Blue), (**B**) KL-S-1D2 (pink), (**C**) KL-S-3A7 (green) and (**D**) 1F7 (orange) bound to SARS-CoV-2 stabilized trimer (grey). A model of spike trimer bound to ACE2 receptor (PDB# 7KNB) is fit into each density to illustrate their potential for blocking receptor binding.

## Discussion

The RBD of the spike protein of SARS-CoV-2 is relatively plastic and can tolerate extensive mutations, at least in *vitro*. The plasticity of the RBD is alarming because extensive changes in the RBD could reduce the efficacy of current vaccines and additional booster vaccinations with updated vaccines may be needed for protection in the future (15, 16). We tested all 14 isolated mAbs for binding to a whole panel of mutant RBDs. While some mAbs lost binding for many mutant RBDs, other mAbs maintained binding well across the board. However, binding was not in all cases directly linked to neutralization. All of the neutralizing mAbs maintained binding and neutralizing activity to B.1.1.7 (N501Y) relatively well. However, two mAbs lost neutralizing activity against B.1.351 and one of these mAbs only showed a relatively low reduction in binding to E484K and B.1.351 RBDs. The second one showed a stronger reduction in binding which agrees better with loss of neutralizing activity. Other hotspots for loss of binding for neutralizing antibodies included amino acid positions 487 and 490.

For four of the neutralizing mAbs low resolution structures were solved using single particle EM. They included KL-S-1D2 which lost neutralizing activity to B.1.351. Our low resolution structural analysis precludes interpretation of molecular interactions but the reduction or loss of neutralization of B1.1.7 and B.1.351 by KL-S-1D2 suggests that N501 and E484 form critical interactions.

Protection *in vivo* by neutralizing mAbs could be a function of Fc-Fc receptor interaction. This has been shown for other mAbs developed against SARS-CoV-2 which showed less protection *in vivo* when the Fc was mutated (25). While the role of Fc-FcR interactions based effector functions for SARS-CoV-2 targeting antibodies is not fully understood yet, it is likely that these effector functions contribute to protection (26). This has also been demonstrated for influenza viruses as well as ebolaviruses (27, 28). We tested all isolated mAbs for their protective effect in a mouse model and found that the only correlation with protection was neutralizing activity while non-neutralizing antibodies had no effect. However, there is an important caveat that needs to be discussed for this experiment. All non-neutralizing antibodies that we isolated were of the IgG1 subtype, which in mice, is known to have low affinity for activating FcRs. This is in contrast to murine IgG2a and IgG2b which have high affinity for these FcRs. Therefore, we can only conclude that non-Fc-FcR based interactions do not contribute to protection by non-neutralizing antibodies. In fact, the two antibodies that provided the best protection, especially on day 3, KL-S-1D2 and KL-S-2C3, are both of the IgG2a subtype. While KL-S-1D2 showed the best *in vitro* neutralization of all isolated mAbs, which could cause this phenotype, KL-S-2C3’s *in vitro* activity was lower but still showed stronger activity *in vivo* than other mAbs. This could be seen as evidence that Fc-FcR interactions, especially engagement with activating FcRs, which is an important component of protection. Of note, the vast majority of antibodies induced in humans to SARS-CoV-2 spike by natural infection or vaccination are IgG1 and in humans – unlike in mice - IgG1 has strong affinity for activating FcRs (29).

In summary, we describe several antibodies to the SARS-CoV-2 RBD that maintain strong neutralizing activity against the B.1.1.7 as well as B.1.351 variant. These mAbs, if humanized, may be further developed into ‘variant resistant’ therapeutics.

## Acknowledgements

We would like to thank Dr. Randy Albrecht for oversight of the conventional Biosafety Level 3 biocontainment facility which made this work possible. This work was partially funded by the NIAID Collaborative Influenza Vaccine Innovation Centers (CIVIC) contract 75N93019C00051, NIAID Center of Excellence for Influenza Research and Surveillance (CEIRS, contract # HHSN272201400008C), by the generous support of the JPB Foundation and the Open Philanthropy Project (research grant 2020-215611 (5384); and by anonymous donors.

## Author contributions

All authors have reviewed and approved the manuscript. F.A designed and performed experiments, analyzed data, and drafted the manuscript. S.S performed animal experiments. W.L and S.B performed structural analysis and prepared figures. L.C provided virus stocks and analyzed data. A.W supervised the structural work and interpreted data. F.K conceptualized the whole study and drafted the manuscript.

## Conflict of interest statement

The Icahn School of Medicine at Mount Sinai has filed patent applications relating to SARS-CoV-2 serological assays and NDV-based SARS-CoV-2 vaccines which list Florian Krammer as co-inventor. Fatima Amanat is also listed on the serological assay patent application as a co-inventor. Mount Sinai has spun out a company, Kantaro, to market serological tests for SARS-CoV-2. Florian Krammer has consulted for Merck and Pfizer (before 2020), and is currently consulting for Pfizer, Seqirus and Avimex. The Krammer laboratory is also collaborating with Pfizer on animal models of SARS-CoV-2.

## Materials and Methods

### Cells and viruses

Vero.E6 cells (ATCC CRL-1586) were maintained in Dulbecco’s modified Eagle’s medium (DMEM; Life Technologies) which was supplemented with 10% fetal bovine serum (FBS; Corning) as well as antibiotics (100 units/ml penicillin–100μg/ml streptomycin [Pen-Strep; Gibco]), and buffer solution [1 M 4-(2-hydroxyethyl)-1-piperazineethanesulfonic acid (HEPES); Gibco]. SARS-CoV-2 was exclusively handled in a BSL3 facility and passaged in Vero.E6 for three days and the supernatant from infected cells was clarified via centrifugation at 1000 g for 5 mins. The virus stock was titered in Vero.E6 cells as well.

### Generation of monoclonal antibodies

All animal work was performed by adhering to institutional regulation as well as Institutional Animal Care and Use Committee (IACUC) guidelines. Six to eight weeks old, female, mice (Jackson Laboratories) were immunized with 3 ugs of purified RBD of SARS-CoV-2 mixed with 10 ugs of poly I:C (Invivogen) twice with 3 weeks interval (17, 29). All immunizations were administered via the intramuscular route. Finally, mice were immunized again with 100 ugs of RBD along with 10 ugs of poly I:C. Three days later, the mouse was sacrificed and the spleen was extracted in a sterile manner. The splenocytes were washed with phosphate buffered saline (Life Technologies; PBS) and then fused with SP2/o myeloma cells (ATCC) using polyethylene glycol (Sigma-Aldrich catalog # P7181). Hybridoma supernatants were screened in an enzyme-linked immunosorbent assay (ELISA) assay as described in the below section. Desirable hybridomas that secreted IgG were selected and expanded. Supernatants was collected at the end, filtered using a 0.22 um filter and then purified via Protein G Sepharose (GE Health) using gravity flow (18, 27, 28, 30–34).

### ELISA

Ninety-six well, flat-bottom, Immulon 4 HBX plates (Thermo Scientific) were coated overnight at 4°C with 50 uls/well of a 2 ug/ml solution of each respective protein in PBS. The next day, coating solution was discarded. One hundred uls per well of 3% non-fat milk prepared in PBS containing 0.1% of Tween-20 (Fisher Bioreagents; T-PBS) was added to the plates for one hour at room temperature (RT) to block the plates. Antibody dilutions were prepared in 1% non-fat milk in T-PBS. The starting concentration used for each antibody was 30 ug/ml and three-fold dilutions were subsequently prepared. After the blocking solution had been on the plates for 1 hour, the antibody dilutions were added for 1 hour at RT. Next, the plates were washed thrice with 250 uls per well of T-PBS. The secondary solution was also prepared in 1% non-fat milk in T-PBS. For mouse antibodies, anti-mouse IgG conjugated to horseradish peroxidase (Rockland) was used at a dilution of 1:3000. For human antibodies, anti-human IgG Fab was used at the same dilution. After 1 hour, plates were washed thrice with 250 uls per well of T-PBS and developing solution was prepared using SigmaFast o-phenylenediamine dihydrochloride (OPD). One hundred uls per well of developing solution was added for exactly ten minutes after which the reaction was stopped by addition of 50 uls per well of 3M HCl (Fisher Bioreagents). Optical density at 490 nanometers was measured using a plate reader, BioTek Synergy H1. All data was analyzed using GraphPad Prism 7. An anti-histidine antibody was used for ELISAs as positive control (Takara, catalog # 631212).

### Microneutralization assay

All antibodies were tested for neutralization capability in a neutralization assay with authentic SARS-CoV-2 isolate USA-WA1/2020 (BEI Resources NR-52281), isolate hCoV-19/South Africa/KRISP-K005325/2020 (BEI Resources NR-54009), and hCoV-19/England/204820464/2020 (BEI Resources NR-54000) in the BSL-3 facility. All viruses were obtained from BEI resources and propagated in Vero.E6 cells. Twenty-thousand Vero.E6 cells were seeded in a 96-well cell culture plate and used the next day. Antibody dilutions were prepared starting at 30 ug/ml and 3-fold subsequent dilutions were prepared. The protocol has been described earlier (21, 29, 35, 36). Cells were stained for the nucleoprotein and quantified. Percent inhibition was calculated and IC_50_s were obtained (35). All viruses were subjected to deep sequencing to ensure that no mutations had taken place in culture. The polybasic cleavage site changed to WRAR in the B.1.351 during passage in cell culture (as known for this virus at BEI Resources) and no other unexpected mutations occurred.

### *In vivo* mouse challenge studies

All work with SARS-CoV-2 was performed in a BSL-3 facility. Six -to 8-weeks old, female, BALB/c mice (Jackson Laboratories) were administered an adenovirus containing human ACE2 (AdV-hACE2) via the intranasal route at 2.5×10^8^ plaque forming units (PFUs) per mouse in a final volume of 50 uls. Five days later, each respective antibody was administered via the intraperitoneal route at 10 mg/kg in 100 uls volume. Two hours later, mice were infected with 10^5^ PFUs of SARS-CoV-2 intranasally. Mice were humanely sacrificed on day 3 and day 5 to assess viral titer in the lungs. Lungs were homogenized using a BeadBlaster 24 (Benchmark) homogenizer. Each lung homogenate was tested in a classical plaque assay as described earlier (27, 30).

### Plaque assay

To assess viral titer in the lungs, each homogenate was diluted in 1X minimal essential medium (10X MEM; Life Technologies) supplemented with glutamine, 35% bovine serum albumin (BSA; MP Biomedicals), antibiotics, and HEPES as described earlier (29, 37). Three-hundred thousand Vero.E6 cells were seeded per well in a 12-well cell culture plate and used the next day when the cells were approximately 90% confluent. Media was removed from cells and dilutions were added to the plates and incubated at 37 degree C incubator for 1 hour. Next, the virus dilutions were removed, and cells were overlaid with 2% oxoid agar mixed with 2X MEM. After 3 days, cells were fixed with 10% paraformaldehyde (Polysciences) for 24 hours and then stained with anti-spike antibodies and plaques were counted.

### Histology and immunohistochemistry (IHC)

Mice were administered anesthesia and euthanized by exsanguination of the femoral artery. Lungs were inflated and flushed with 10% formaldehyde by injecting a needle through the trachea on day 4 post infection with SARS-CoV-2. Fixed lung samples were sent for processing to a commercial company, Histowiz. Sections were analyzed, images were taken, and sections were also scored by a pathologist. Scores were assigned by the pathologist based on six parameters, as mentioned in the results section. Both H&E staining as well as IHC was performed. An anti-SARS-CoV nucleoprotein antibody (Novus Biologicals cat. NB100–56576) was used for IHC.

### Expression and purification of recombinant spike proteins for electron microscopy

The SARS-CoV-2 spike construct used for EM studies contains the mammalian-codon-optimized gene encoding residues 1-1208 of the spike followed by a C-terminal T4 fibritin trimerization domain, a HRV3C cleavage site and an 8x-His tag subcloned into the eukaryotic-expression vector pcDNA3.4. Amino acid mutations were introduced in the S1/S2 cleavage site (RRAR to GSAS) along with other stabilizing mutations including the HexaPro mutations (38). The spike trimers were expressed and purified as described previously (39).

### Negative stain EM sample preparation and data collection

Spike protein was complexed with purified Fab at three times molar excess per trimer and incubated for thirty minutes at room temperature. Complexes were diluted to 0.02mg/ml in TBS and 3μl applied to a 400mesh Cu grid, blotted with filter paper, and stained with 2% uranyl formate for 30 seconds. Images were collected on a Tecnai Spirit microscope operating at 120 kV with a FEI Eagle CCD (4k) camera. Particles were picked using DogPicker and 3D classification was done using Relion 3.0 (40, 41).

---

**Supplementary figure 1.** Clinical scores from the lung sections from each antibody group (n=2) are shown. All sections were scored by an independent veterinary pathologist. In addition to negative control, one control group received only AdV-hACE2 while naïve mice were also included as mock control. Scale bar represents 500 um.

## References

1. Zhu N, Zhang D, Wang W, Li X, Yang B, Song J, Zhao X, Huang B, Shi W, Lu R, Niu P, Zhan F, Ma X, Wang D, Xu W, Wu G, Gao GF, Tan W, China Novel Coronavirus I, Research T. 2020. A Novel Coronavirus from Patients with Pneumonia in China, 2019. N Engl J Med 382:727–733.

2. Huang C, Wang Y, Li X, Ren L, Zhao J, Hu Y, Zhang L, Fan G, Xu J, Gu X, Cheng Z, Yu T, Xia J, Wei Y, Wu W, Xie X, Yin W, Li H, Liu M, Xiao Y, Gao H, Guo L, Xie J, Wang G, Jiang R, Gao Z, Jin Q, Wang J, Cao B. 2020. Clinical features of patients infected with 2019 novel coronavirus in Wuhan, China. Lancet 395:497–506.

3. White KM, Rosales R, Yildiz S, Kehrer T, Miorin L, Moreno E, Jangra S, Uccellini MB, Rathnasinghe R, Coughlan L, Martinez-Romero C, Batra J, Rojc A, Bouhaddou M, Fabius JM, Obernier K, Dejosez M, Guillen MJ, Losada A, Aviles P, Schotsaert M, Zwaka T, Vignuzzi M, Shokat KM, Krogan NJ, Garcia-Sastre A. 2021. Plitidepsin has potent preclinical efficacy against SARS-CoV-2 by targeting the host protein eEF1A. Science 371:926–931.

4. Hattori SI, Higashi-Kuwata N, Hayashi H, Allu SR, Raghavaiah J, Bulut H, Das D, Anson BJ, Lendy EK, Takamatsu Y, Takamune N, Kishimoto N, Murayama K, Hasegawa K, Li M, Davis DA, Kodama EN, Yarchoan R, Wlodawer A, Misumi S, Mesecar AD, Ghosh AK, Mitsuya H. 2021. A small molecule compound with an indole moiety inhibits the main protease of SARS-CoV-2 and blocks virus replication. Nat Commun 12:668.

5. Chen P, Nirula A, Heller B, Gottlieb RL, Boscia J, Morris J, Huhn G, Cardona J, Mocherla B, Stosor V, Shawa I, Adams AC, Van Naarden J, Custer KL, Shen L, Durante M, Oakley G, Schade AE, Sabo J, Patel DR, Klekotka P, Skovronsky DM, Investigators B-. 2021. SARS-CoV-2 Neutralizing Antibody LY-CoV555 in Outpatients with Covid-19. N Engl J Med 384:229–237.

6. Weinreich DM, Sivapalasingam S, Norton T, Ali S, Gao H, Bhore R, Musser BJ, Soo Y, Rofail D, Im J, Perry C, Pan C, Hosain R, Mahmood A, Davis JD, Turner KC, Hooper AT, Hamilton JD, Baum A, Kyratsous CA, Kim Y, Cook A, Kampman W, Kohli A, Sachdeva Y, Graber X, Kowal B, DiCioccio T, Stahl N, Lipsich L, Braunstein N, Herman G, Yancopoulos GD, Trial I. 2021. REGN-COV2, a Neutralizing Antibody Cocktail, in Outpatients with Covid-19. N Engl J Med 384:238–251.

7. Letko M, Marzi A, Munster V. 2020. Functional assessment of cell entry and receptor usage for SARS-CoV-2 and other lineage B betacoronaviruses. Nat Microbiol 5:562–569.

8. Hoffmann M, Kleine-Weber H, Schroeder S, Kruger N, Herrler T, Erichsen S, Schiergens TS, Herrler G, Wu NH, Nitsche A, Muller MA, Drosten C, Pohlmann S. 2020. SARS-CoV-2 Cell Entry Depends on ACE2 and TMPRSS2 and Is Blocked by a Clinically Proven Protease Inhibitor. Cell 181:271–280 e8.

9. Wrapp D, Wang N, Corbett KS, Goldsmith JA, Hsieh CL, Abiona O, Graham BS, McLellan JS. 2020. Cryo-EM structure of the 2019-nCoV spike in the prefusion conformation. Science 367:1260–1263.

10. Wajnberg A, Amanat F, Firpo A, Altman DR, Bailey MJ, Mansour M, McMahon M, Meade P, Mendu DR, Muellers K, Stadlbauer D, Stone K, Strohmeier S, Simon V, Aberg J, Reich DL, Krammer F, Cordon-Cardo C. 2020. Robust neutralizing antibodies to SARS-CoV-2 infection persist for months. Science 370:1227–1230.

11. Jackson LA, Anderson EJ, Rouphael NG, Roberts PC, Makhene M, Coler RN, McCullough MP, Chappell JD, Denison MR, Stevens LJ, Pruijssers AJ, McDermott A, Flach B, Doria-Rose NA, Corbett KS, Morabito KM, O’Dell S, Schmidt SD, Swanson PA, 2nd, Padilla M, Mascola JR, Neuzil KM, Bennett H, Sun W, Peters E, Makowski M, Albert J, Cross K, Buchanan W, Pikaart-Tautges R, Ledgerwood JE, Graham BS, Beigel JH, m RNASG. 2020. An mRNA Vaccine against SARS-CoV-2 - Preliminary Report. N Engl J Med 383:1920–1931.

12. Voysey M, Costa Clemens SA, Madhi SA, Weckx LY, Folegatti PM, Aley PK, Angus B, Baillie VL, Barnabas SL, Bhorat QE, Bibi S, Briner C, Cicconi P, Clutterbuck EA, Collins AM, Cutland CL, Darton TC, Dheda K, Dold C, Duncan CJA, Emary KRW, Ewer KJ, Flaxman A, Fairlie L, Faust SN, Feng S, Ferreira DM, Finn A, Galiza E, Goodman AL, Green CM, Green CA, Greenland M, Hill C, Hill HC, Hirsch I, Izu A, Jenkin D, Joe CCD, Kerridge S, Koen A, Kwatra G, Lazarus R, Libri V, Lillie PJ, Marchevsky NG, Marshall RP, Mendes AVA, Milan EP, Minassian AM, et al. 2021. Single-dose administration and the influence of the timing of the booster dose on immunogenicity and efficacy of ChAdOx1 nCoV-19 (AZD1222) vaccine: a pooled analysis of four randomised trials. Lancet.

13. Chen AT, Altschuler K, Zhan SH, Chan YA, Deverman BE. 2021. COVID-19 CG enables SARS-CoV-2 mutation and lineage tracking by locations and dates of interest. Elife 10.

14. McCallum M, Marco A, Lempp F, Tortorici MA, Pinto D, Walls AC, Beltramello M, Chen A, Liu Z, Zatta F, Zepeda S, di Iulio J, Bowen JE, Montiel-Ruiz M, Zhou J, Rosen LE, Bianchi S, Guarino B, Fregni CS, Abdelnabi R, Caroline Foo SY, Rothlauf PW, Bloyet LM, Benigni F, Cameroni E, Neyts J, Riva A, Snell G, Telenti A, Whelan SPJ, Virgin HW, Corti D, Pizzuto MS, Veesler D. 2021. N-terminal domain antigenic mapping reveals a site of vulnerability for SARS-CoV-2. bioRxiv.

15. Greaney AJ, Starr TN, Gilchuk P, Zost SJ, Binshtein E, Loes AN, Hilton SK, Huddleston J, Eguia R, Crawford KHD, Dingens AS, Nargi RS, Sutton RE, Suryadevara N, Rothlauf PW, Liu Z, Whelan SPJ, Carnahan RH, Crowe JE, Jr., Bloom JD. 2021. Complete Mapping of Mutations to the SARS-CoV-2 Spike Receptor-Binding Domain that Escape Antibody Recognition. Cell Host Microbe 29:44–57 e9.

16. Starr TN, Greaney AJ, Addetia A, Hannon WW, Choudhary MC, Dingens AS, Li JZ, Bloom JD. 2021. Prospective mapping of viral mutations that escape antibodies used to treat COVID-19. Science 371:850–854.

17. Tan GS, Lee PS, Hoffman RM, Mazel-Sanchez B, Krammer F, Leon PE, Ward AB, Wilson IA, Palese P. 2014. Characterization of a broadly neutralizing monoclonal antibody that targets the fusion domain of group 2 influenza A virus hemagglutinin. J Virol 88:13580–92.

18. Wohlbold TJ, Chromikova V, Tan GS, Meade P, Amanat F, Comella P, Hirsh A, Krammer F. 2016. Hemagglutinin Stalk- and Neuraminidase-Specific Monoclonal Antibodies Protect against Lethal H10N8 Influenza Virus Infection in Mice. J Virol 90:851–61.

19. Amanat F, Meade P, Strohmeier S, Krammer F. 2019. Cross-reactive antibodies binding to H4 hemagglutinin protect against a lethal H4N6 influenza virus challenge in the mouse model. Emerg Microbes Infect 8:155–168.

20. Rathnasinghe R, Strohmeier S, Amanat F, Gillespie VL, Krammer F, Garcia-Sastre A, Coughlan L, Schotsaert M, Uccellini MB. 2020. Comparison of transgenic and adenovirus hACE2 mouse models for SARS-CoV-2 infection. Emerg Microbes Infect 9:2433–2445.

21. Amanat F, Strohmeier S, Rathnasinghe R, Schotsaert M, Coughlan L, Garcia-Sastre A, Krammer F. 2020. Introduction of two prolines and removal of the polybasic cleavage site leads to optimal efficacy of a recombinant spike based SARS-CoV-2 vaccine in the mouse model. bioRxiv.

22. Tseng CT, Sbrana E, Iwata-Yoshikawa N, Newman PC, Garron T, Atmar RL, Peters CJ, Couch RB. 2012. Immunization with SARS coronavirus vaccines leads to pulmonary immunopathology on challenge with the SARS virus. PLoS One 7:e35421.

23. Barnes CO, Jette CA, Abernathy ME, Dam KA, Esswein SR, Gristick HB, Malyutin AG, Sharaf NG, Huey-Tubman KE, Lee YE, Robbiani DF, Nussenzweig MC, West AP, Jr., Bjorkman PJ. 2020. SARS-CoV-2 neutralizing antibody structures inform therapeutic strategies. Nature 588:682–687.

24. Pinto D, Park YJ, Beltramello M, Walls AC, Tortorici MA, Bianchi S, Jaconi S, Culap K, Zatta F, De Marco A, Peter A, Guarino B, Spreafico R, Cameroni E, Case JB, Chen RE, Havenar-Daughton C, Snell G, Telenti A, Virgin HW, Lanzavecchia A, Diamond MS, Fink K, Veesler D, Corti D. 2020. Cross-neutralization of SARS-CoV-2 by a human monoclonal SARS-CoV antibody. Nature 583:290–295.

25. Schafer A, Muecksch F, Lorenzi JCC, Leist SR, Cipolla M, Bournazos S, Schmidt F, Maison RM, Gazumyan A, Martinez DR, Baric RS, Robbiani DF, Hatziioannou T, Ravetch JV, Bieniasz PD, Bowen RA, Nussenzweig MC, Sheahan TP. 2021. Antibody potency, effector function, and combinations in protection and therapy for SARS-CoV-2 infection in vivo. J Exp Med 218.

26. Gorman MJ, Patel N, Guebre-Xabier M, Zhu A, Atyeo C, Pullen KM, Loos C, Goez-Gazi Y, Carrion R, Tian J-H, Yaun D, Bowman K, Zhou B, Maciejewski S, McGrath ME, Logue J, Frieman MB, Montefiori D, Mann C, Schendel S, Amanat F, Krammer F, Saphire EO, Lauffenburger D, Greene AM, Portnoff AD, Massare MJ, Ellingsworth L, Glenn G, Smith G, Alter G. 2021. Collaboration between the Fab and Fc contribute to maximal protection against SARS-CoV-2 in nonhuman primates following NVX-CoV2373 subunit vaccine with Matrix-M™ vaccination. bioRxiv:2021.02.05.429759.

27. Duehr J, Wohlbold TJ, Oestereich L, Chromikova V, Amanat F, Rajendran M, Gomez-Medina S, Mena I, tenOever BR, Garcia-Sastre A, Basler CF, Munoz-Fontela C, Krammer F. 2017. Novel Cross-Reactive Monoclonal Antibodies against Ebolavirus Glycoproteins Show Protection in a Murine Challenge Model. J Virol 91.

28. Asthagiri Arunkumar G, Ioannou A, Wohlbold TJ, Meade P, Aslam S, Amanat F, Ayllon J, Garcia-Sastre A, Krammer F. 2019. Broadly Cross-Reactive, Nonneutralizing Antibodies against Influenza B Virus Hemagglutinin Demonstrate Effector Function-Dependent Protection against Lethal Viral Challenge in Mice. J Virol 93.

29. Amanat F, Stadlbauer D, Strohmeier S, Nguyen THO, Chromikova V, McMahon M, Jiang K, Arunkumar GA, Jurczyszak D, Polanco J, Bermudez-Gonzalez M, Kleiner G, Aydillo T, Miorin L, Fierer DS, Lugo LA, Kojic EM, Stoever J, Liu STH, Cunningham-Rundles C, Felgner PL, Moran T, Garcia-Sastre A, Caplivski D, Cheng AC, Kedzierska K, Vapalahti O, Hepojoki JM, Simon V, Krammer F. 2020. A serological assay to detect SARS-CoV-2 seroconversion in humans. Nat Med 26:1033–1036.

30. Wohlbold TJ, Podolsky KA, Chromikova V, Kirkpatrick E, Falconieri V, Meade P, Amanat F, Tan J, tenOever BR, Tan GS, Subramaniam S, Palese P, Krammer F. 2017. Broadly protective murine monoclonal antibodies against influenza B virus target highly conserved neuraminidase epitopes. Nat Microbiol 2:1415–1424.

31. Amanat F, Duehr J, Oestereich L, Hastie KM, Ollmann Saphire E, Krammer F. 2018. Antibodies to the Glycoprotein GP2 Subunit Cross-React between Old and New World Arenaviruses. mSphere 3.

32. Stadlbauer D, Amanat F, Strohmeier S, Nachbagauer R, Krammer F. 2018. Cross-reactive mouse monoclonal antibodies raised against the hemagglutinin of A/Shanghai/1/2013 (H7N9) protect against novel H7 virus isolates in the mouse model. Emerg Microbes Infect 7:110.

33. Stadlbauer D, Rajabhathor A, Amanat F, Kaplan D, Masud A, Treanor JJ, Izikson R, Cox MM, Nachbagauer R, Krammer F. 2017. Vaccination with a Recombinant H7 Hemagglutinin-Based Influenza Virus Vaccine Induces Broadly Reactive Antibodies in Humans. mSphere 2.

34. Strohmeier S, Amanat F, Krammer F. 2019. Cross-Reactive Antibodies Binding to the Influenza Virus Subtype H11 Hemagglutinin. Pathogens 8.

35. Amanat F, White KM, Miorin L, Strohmeier S, McMahon M, Meade P, Liu WC, Albrecht RA, Simon V, Martinez-Sobrido L, Moran T, Garcia-Sastre A, Krammer F. 2020. An In Vitro Microneutralization Assay for SARS-CoV-2 Serology and Drug Screening. Curr Protoc Microbiol 58:e108.

36. Sun W, Leist SR, McCroskery S, Liu Y, Slamanig S, Oliva J, Amanat F, Schafer A, Dinnon KH, 3rd, Garcia-Sastre A, Krammer F, Baric RS, Palese P. 2020. Newcastle disease virus (NDV) expressing the spike protein of SARS-CoV-2 as a live virus vaccine candidate. EBioMedicine 62:103132.

37. Duehr J, McMahon M, Williamson B, Amanat F, Durbin A, Hawman DW, Noack D, Uhl S, Tan GS, Feldmann H, Krammer F. 2020. Neutralizing Monoclonal Antibodies against the Gn and the Gc of the Andes Virus Glycoprotein Spike Complex Protect from Virus Challenge in a Preclinical Hamster Model. mBio 11.

38. Hsieh CL, Goldsmith JA, Schaub JM, DiVenere AM, Kuo HC, Javanmardi K, Le KC, Wrapp D, Lee AG, Liu Y, Chou CW, Byrne PO, Hjorth CK, Johnson NV, Ludes-Meyers J, Nguyen AW, Park J, Wang N, Amengor D, Lavinder JJ, Ippolito GC, Maynard JA, Finkelstein IJ, McLellan JS. 2020. Structure-based design of prefusion-stabilized SARS-CoV-2 spikes. Science 369:1501–1505.

39. Brouwer PJM, Caniels TG, van der Straten K, Snitselaar JL, Aldon Y, Bangaru S, Torres JL, Okba NMA, Claireaux M, Kerster G, Bentlage AEH, van Haaren MM, Guerra D, Burger JA, Schermer EE, Verheul KD, van der Velde N, van der Kooi A, van Schooten J, van Breemen MJ, Bijl TPL, Sliepen K, Aartse A, Derking R, Bontjer I, Kootstra NA, Wiersinga WJ, Vidarsson G, Haagmans BL, Ward AB, de Bree GJ, Sanders RW, van Gils MJ. 2020. Potent neutralizing antibodies from COVID-19 patients define multiple targets of vulnerability. Science 369:643–650.

40. Voss NR, Yoshioka CK, Radermacher M, Potter CS, Carragher B. 2009. DoG Picker and TiltPicker: software tools to facilitate particle selection in single particle electron microscopy. J Struct Biol 166:205–13.

41. Zivanov J, Nakane T, Forsberg BO, Kimanius D, Hagen WJ, Lindahl E, Scheres SH. 2018. New tools for automated high-resolution cryo-EM structure determination in RELION-3. Elife 7.

